# Increasing central serotonin with 5-HTP disrupts the inhibition of social gaze in non-human primates

**DOI:** 10.1101/2021.02.26.431901

**Authors:** Hannah Weinberg-Wolf, Nicholas A. Fagan, Olga Dal Monte, Steve W. C. Chang

## Abstract

To competently navigate the world, individuals must flexibly balance distinct aspects of social gaze, orienting toward others and inhibiting orienting responses, depending on the context. These behaviors are often disrupted in patient populations treated with serotonergic drugs. However, the field lacks a clear understanding of how the serotonergic system mediates social orienting and inhibiting behaviors. Here, we tested how increasing central concentrations of serotonin with the direct precursor 5-Hydroxytryptophan (5-HTP) would modulate the ability of rhesus macaques to use eye movements to flexibly orient to, or inhibit orienting to, faces. Systemic administrations of 5-HTP effectively increased central serotonin levels and impaired flexible orientation and inhibition. Critically, 5-HTP selectively impaired the ability of monkeys to inhibit orienting to face images, whereas it similarly impaired orienting to face and control images. 5-HTP also caused monkeys to perseverate on their gaze responses, making them worse at flexibly switching between orientating and inhibiting behaviors. Furthermore, the effects of 5-HTP on performance correlated with a constriction of the pupil, an increased time to initiate trials, and an increased reaction time, suggesting that the disruptive effects of 5-HTP on social gaze behaviors are likely driven by a downregulation of arousal and motivational states. Taken together, these findings provide causal evidence for a modulatory relationship between 5-HTP and social gaze behaviors in non-human primates and offer translational insights for the role of the serotonergic system in social gaze.

## Introduction

Serotonergic agents are broadly prescribed to individuals suffering from numerous psychiatric disorders including depression, anxiety disorders, personality disorders, and social disorders including autism spectrum disorder (ASD) (Abi-Dargham et al., 1997; Dell’Osso et al., 2010; Williams et al., 2011; Cowen and Browning, 2015; Healy, 2015; Muller et al., 2016). However, outcomes of serotonergic interventions are variable and difficult to predict (Shaw et al., 2002; Turner et al., 2006; Chekroud et al., 2016). To better understand serotonergic interventions, and to improve treatment outcomes, it is important to conduct controlled, causal, experiments which test central serotonin’s effect on cognitive functions.

Competently navigating the world requires balancing the acts of orienting toward environmental stimuli and inhibiting orienting responses. This behavioral regulation demands coordination among the neural systems underlying motivation, inhibition, attention, and flexibility (Elliot, 2006; Carver et al., 2008; Weinberg-Wolf and Chang, 2019; Roberts et al., 2020). Among environmental stimuli, faces are particularly common and important for humans and non-human primates. Many species of primates, including humans, live in large, complex, societies and therefore orient to others’ faces to learn from conspecifics and to opportunistically gain information about fecundity and social status. However, many primate species must also inhibit orienting to faces to avoid accidentally aggressing to conspecifics and to avoid losing opportunities to gain non-social information about food sources and environmental risks (Weinberg-Wolf and Chang, 2019).

Clinically, disruptions in social orienting are frequently observed in ASD and Williams Syndrome (WS). Infants with ASD lack early social predispositions (Klin et al., 2002; Dawson et al., 2004) and these deficits continue into adulthood (Pelphrey et al., 2002; Kliemann et al., 2010). Evidence further suggests that the deliberate recognition of, and orienting toward, different emotional expressions are impaired in ASD (Sigman et al., 1992; Bacon et al., 1998; Humphreys et al., 2007). On the other hand, individuals with WS, a rare genetic disorder associated with hyper-sociality, exhibit heightened social engagement, increased attention to faces, and uninhibited approach to strangers (Barak and Feng, 2016). Central serotonergic functions are suspected to be dysregulated in both ASD and WS (August and Realmuto, 1989; Williams et al., 2011; Barak and Feng, 2016; Muller et al., 2016; Fan et al., 2020; Lew et al., 2020).

The serotonergic system is implicated in diverse aspects of behavioral inhibition (Roberts et al., 2020). Decreasing central serotonin disrupts impulse control in humans (LeMarquand et al., 1999; Cools et al., 2005; Rubia et al., 2005; Crockett et al., 2010), non-human primates (Clarke et al., 2004; Clarke et al., 2005), and rodents (Harrison et al., 1997b, 1999; Matias et al., 2017; Bailey et al., 2018; Lottem et al., 2018). Importantly, serotonin is also associated with flexibility; disrupting serotonergic function impacts reversal learning and perseverance during choice tasks (Roberts et al., 2020). Modulating serotonin function can also alter cognitive biases, attention, and motivation, with these effects often being strongest within the social domain (Young, 1996; Riedel, 2004; Merens et al., 2007; Mendelsohn et al., 2009; Silber and Schmitt, 2010; Roberts et al., 2020). These findings indicate that the serotonergic system is well positioned to affect flexible social orienting behaviors.

Causal manipulations of serotonergic function on behavior, especially repeated within-subject studies, are rare in non-human primates. We have previously shown that acute delivery of the direct serotonin precursor 5-Hydroxytroptohan (5-HTP) increases central concentrations of macaques’ serotonin with concomitant changes attention to faces and landscapes (Weinberg-Wolf et al., 2018). Despite growing evidence of serotonin’s role in inhibition, it remains unclear how increasing central serotonin availability affects social gaze inhibition along with the flexible balancing of orienting and inhibiting gaze. Causal characterizations of such behavioral changes in non-human primate models, in which these behaviors are understudied, will help support continued translational efforts. Here, we utilized a rhesus macaque model to examine how increasing central serotonin with 5-HTP modulates social gaze behaviors when monkeys were cued to either orient towards faces or inhibit this orienting response.

## Materials and Methods

### Animals

Three adult (2 male and 1 female; monkeys E, H, T; age 7 years) rhesus monkeys (*Macaca mulatta*) served as subjects. Subjects weighed between 10 and 15kg throughout the duration of the study. Subjects were housed single (n=2/3) or with a single pair (n=1/3), kept on a 12-hr light/dark cycle, had unrestricted access to food 24 hours a day, and had controlled access to fluid during testing. All procedures were reviewed and approved by the Yale University Institutional Animal Care and Use Committee.

### Surgery

All subjects received a surgically implanted head-restraining prosthesis (Grey Matter Research) to allow for accurate tracking of eye movements. At the time of surgery, anesthesia was induced with ketamine hydrochloride (10mg/kg) and maintained with isoflurane (1.0–3.0%, to effect). During surgery, subjects received isotonic fluids via an intravenous drip, and aseptic procedures were employed. Heart rate, respiration rate, blood pressure, expired CO_2_, and body temperature were monitored throughout the procedure. After the head restraining device implantation was completed, the wound around the base was closed in anatomical layers. Subjects received a peri- and post-operative treatment regimen consisting of 0.01mg/kg buprenorphine every 12 hours for 3 days, 0.1mg/kg meloxicam once daily for 3 days, and 5mg/kg baytril once daily for 10 days. Subjects were allowed 40+ days of recovery after the implant surgery before training began and were slowly acclimated to head restraint over a week of training.

### Pharmacological methods

All pharmacological treatments were administered daily (consistently at either 12:30 or at 16:00 depending on the subject) and intramuscularly (i.m.) exactly one hour before testing. Administered volume was consistently between 3.0 and 5.0ml depending on the weight of the subject. 5-HTP (Sigma) was suspended in sterile water and given at 20mg/kg. Vehicle injections consisted of equal volumes of sterile saline. Each subject received a total of 8 injections of saline and 8 injections of 20mg/kg 5-HTP on strictly alternating days. The dose of 5-HTP was selected on the basis of our previous work (Weinberg-Wolf et al., 2018) indicating that 20mg/kg 5-HTP effectively increases the concentration of 5-HTP and serotonin in cisternal cerebrospinal fluid (CSF). Previous studies in rodents and human subjects (Griebel, 1995; Turner et al., 2006) have indicated that 5-HTP doses less than 20mg/kg do not reliably produce discernable behavioral effects while doses greater than 60mg/kg can inadvertently increase circulating catecholamines by displacing them from storage granules, thereby temporarily enhancing postsynaptic catecholaminergic stimulation (Lichtensteiger et al., 1967).

### Cisternal CSF sample collection and assays

To confirm whether i.m. 5-HTP crossed the blood brain barrier in rhesus macaques, and to test if injections indeed increased central levels of 5-HTP as well as serotonin, we sampled CSF from the cisterna magna from 5 rhesus monkeys (4 male and 1 female; monkeys E, H, T, K, L; aged 5–8 years, 5.5 ± 1.2) after they had received an i.m. injection of saline or 20mg/kg 5-HTP with a minimum of 2 weeks between each CSF collection date. We counterbalanced and randomized the subject order of CSF sampling between saline and 5-HTP. Each CSF draw occurred one-hour post injection, the same time after injection that animals began data collection in the current study.

Cisternal CSF was obtained with cervical punctures, which are preferred over lumbar punctures for accurately tracking concentrations of monoamines and monoamine metabolites in cortical and subcortical structures due to its greater proximity to the brain and its clearance from the spinal space (Anderson et al., 1987b; Anderson et al., 2002; Anderson et al., 2005). Punctures targeted the cisterna magna through the atlanto-occipital membrane. Approximately 1.5ml of CSF was drawn using a 24–27-gauge needle. Monkeys were first anesthetized with ketamine (3mg/kg, i.m.) and dexdomitor (0.075mg/kg, i.m.) and anesthesia was reversed with antisedan (0.075mg/kg, i.m.) once the animal was returned home. CSF was immediately labeled and frozen on dry ice before being transferred to a −80-degree Celsius freezer.

Samples with gross blood contamination (>0.1%), as indicated by pink coloration, were excluded prior to screening for hemoglobin. Limiting blood contamination to <0.1% was sufficient to ensure that analyses other than serotonin were not affected by blood. However, all CSF samples analyzed for serotonin were screened more rigorously for blood contamination by measuring hemoglobin using Multistix 8 SG reagent strips for urinalysis (Bayer Corp., Elkhart, IN), which can detect approximately 0.2μg/ml of hemoglobin. As previously demonstrated, screening for hemoglobin and using only those samples with <10 ppm blood limited blood-derived serotonin in CSF to less than 10pg/ml (Anderson et al., 2005). Neurochemical analyses levels of CSF 5-HTP and serotonin were determined using reverse-phase high performance liquid chromatography (HPLC) as previously described (Anderson et al., 1987a; Anderson et al., 1987b; Anderson et al., 1990; Anderson et al., 2002).

Samples with any detectable blood contamination were excluded from analysis – a total of 2 samples were excluded. The final 5-HTP CSF data therefore included samples collected after injection of saline and 20mg/kg 5-HTP from 4 subjects with an additional subject contributing a sample at saline. The final serotonin CSF data set was more restricted and includes saline data from 4 subjects and 20mg/kg 5-HTP data from 3 subjects. Changes in CSF concentrations of 5-HTP and serotonin were each assessed using a two tailed independent *t*-test.

### Experimental design

Monkeys performed the experimental task while sitting in a primate chair (Precision Engineering Co.) in a testing room and used eye movements to interact with stimuli on an LCD monitor, positioned 36cm away from the subject with a temporal resolution of 2ms. MATLAB (Math Works) with Psychtoolbox (Brainard, 1997) and Eyelinktoolbox (Pelli, 1997) was used to display stimuli and process eye position data. Horizontal and vertical eye positions were sampled at 1,000Hz using an infrared eye monitor camera system (SR Research Eyelink). Monkeys initiated each trial by fixating on a white fixation square (7.3×7.3 visual degrees, deg) at the center for the screen for 150ms. Upon successful fixation, a central instructional cue (7.3×7.3 deg) appeared at the center of the screen and, at the same time, a target image (9.8×9.8 deg) appeared in the right or left periphery of the screen at a 29.3-deg eccentricity. On 50% of pseudo-random trials (orienting trials), the central instructional cue was a red square, directing the subject to saccade to the peripheral target image in order to receive a juice reward (**Fig. 1A**). On the other 50% of pseudo-random trials (inhibition trials), the instructional cue was a blue square, directing subjects to inhibit orienting their gaze towards the target image receive the reward (**Fig. 1A**). From the time of image onset, the subjects had 750ms to orient or inhibit orienting to the image. Images disappeared as soon as subjects’ eyes entered the image (either on correct orientating trials or incorrect inhibition trials) or after the 750ms window on correct inhibition trials. A solenoid valve controlled the delivery of 0.5ml of apple juice per each correct orienting or inhibiting response.

**Figure 1.**
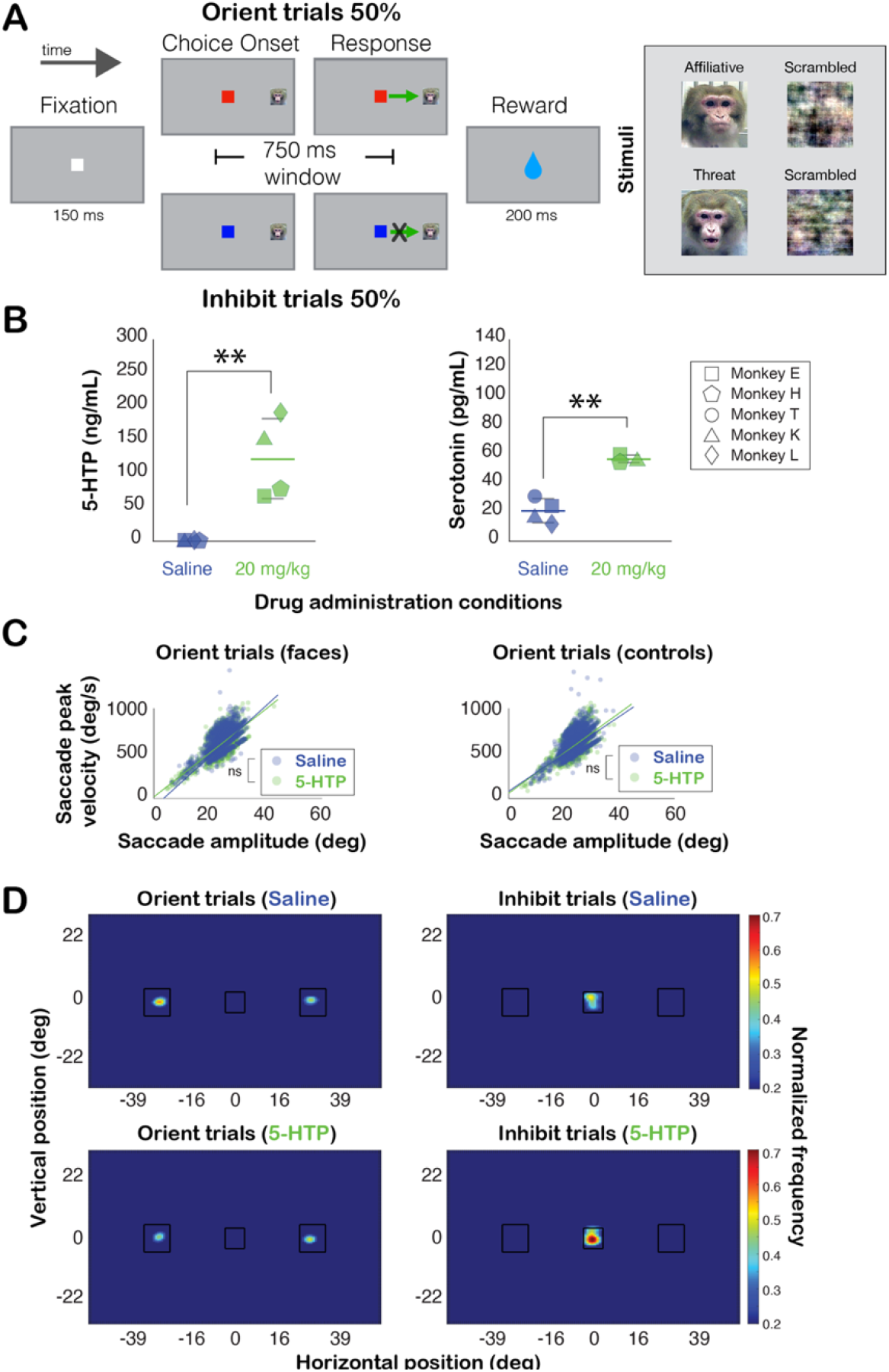
Behavioral task and CSF concentrations of 5-HTP and serotonin following 5-HTP and saline administrations. **A)** Behavioral task sequence. Example face and luminance-matched scrambled control images are shown on the right. The target image appeared either at the right or left of the instructional cue (see Materials and Methods). **B)** (Left) CSF concentration of 5-HTP illustrating the central concentration of 5-HTP after i.m. injection of saline (blue) or 20mg/kg 5-HTP (green). (Right) CSF concentration of serotonin illustrating the central concentration of serotonin after i.m. injection of saline (blue) or 20mg/kg 5-HTP (green). The average CSF concentration per condition is represented by a colored line, and shorter grey lines represent the standard error. Each shape represents individual monkey’s data. **, *P* < 0.01, t-test. **C)** Quantifications of saccade kinematics during successful orient trials to faces (left) and control images (right) during both 5-HTP (blue) and saline (green) sessions. **D)** Fixation density heatmaps (normalized fixation frequency) of correct orient and inhibit trials for 5-HTP and saline conditions. Black outlines represent the stimuli (instructional cue and target images).

The peripheral visual images were unaltered conspecific face stimuli taken from a large library of static monkey face images described by Gothard and colleagues (Gothard et al., 2004). Stimulus monkeys displayed either threatening expression or affiliative lipsmacks with a direct gaze. Subjects had never interacted with any of the monkeys depicted in these images. We divided the total number of images into four unique sets. Each set consisted of 24 conspecific faces per each of the two facial expression categories, resulting in 48 unique faces per set for a total of 192 images. Each image set also contained equal numbers (48 images per set) of luminance-matched scrambled faces (control images). We collected two days of data per each image set for each drug condition. To preclude any order effects, we counterbalanced the order in which we selected image sets while ensuring that subjects were never exposed to the same set of images during two sessions in a row. Within a single session of data collection, we also counterbalanced and randomized the order of image presentation to preclude any order effects.

Initiated trials were defined as those in which the monkeys successfully held gaze fixation for 150ms during the fixation period at the beginning of the trial and thus successfully progressed to view an instructional cue and target image. Upon breaking fixation during this period, the trial was aborted, no cue or image appeared, and the animal was not rewarded. Instead, they received a 1.5sec timeout with a blank screen. Finally, we excluded initiated trials from the analysis that were associated with a pupil size during the fixation window that was more than two standard deviations away from the mean pupil size within drug condition, trial type, and image category. Using this criterion, we only excluded 3% of initiated trials.

### Statistical analysis

To compare saccade kinematics between 5-HTP and saline, we calculated the peak saccade velocity (deg/sec) of orienting saccades (made during the 750-ms window aligned to the onset of the instructional cue and the visual image, **Fig. 1A**) and plotted them as a function of the peak saccade amplitude (deg) for each saccade during 5-HTP and saline sessions, separately for trials with faces and control images. To examine gaze fixation patterns of successfully performed trials, we constructed fixation density maps of the screen, spanning both the instructional cue and the target image locations, for all gaze fixations made from 300ms to 750ms (aligned to the onset of the instructional cue and the visual image), separately for 5-HTP and saline trials. We average how many gaze samples occurred in bounds of the stimulus, separately for the left and right target image and the instructional cue. We then tested for any differences in the number of gaze samples between 5-HTP and saline sessions, separately for orientation and inhibition trials, using Wilcoxon rank sum tests.

Our primary measure of interest was performance in the task. To directly examine whether 5- HTP modulated performance, we calculated the percent of initiated trials that monkeys completed correctly. Percent correct was assessed using analysis of variance (ANOVA) models. The first model specified animal identity (Monkey E, Monkey H, Monkey T), drug (saline versus 20mg/kg 5-HTP), trial type (orientating versus inhibition), image type (face versus scrambled), and image valence (threat versus appetitive) as fixed factors. The second model used data which was normalized relative to saline, as a percent change in performance due to 5-HTP, to control for individual differences between monkeys and specified trial type (orienting versus inhibition), image type (face versus control), and image valence (threat versus appetitive) as fixed factors. Finally, the third model examined performance separately for orienting and inhibition trials, and specified image type (face versus control) and image valence (threat versus appetitive). Direct post hoc comparisons were made with two-tailed independent *t*-tests and *P*-values were corrected for multiple comparisons with Tukey tests. To further investigate the stability of performance over the course of a given session, we plotted performance over time for orienting or inhibiting orienting to face and scrambled images during saline and 5-HTP sessions. Performance was calculated across 240-sec bins with 20-sec steps. Differences in performance between saline and 5-HTP sessions were assessed by conducting a Wilcoxon sign rank test for each of these bins.

To query if 5-HTP impacted monkeys’ flexibility in the current task, we quantified performance (as percent correct) as a function of the following four preceding contexts: 1) the previous trial was correct and the same type (orientation [O] or inhibition [I]) as the current trial (same; O→O or I→I); 2) the previous trial was correct, but it was a different trial type than the current type (switch; O→I or I→O); 3) the previous two trials were both correct and both the same as the current trial (same; O→O→O or I→I→I); and finally 4) the previous two trials were both correct but were both a different trial type than the current trial type (switch; O→O→I or I→I→O). We then controlled for individual differences between monkeys by quantifying the percent change in performance due to 5-HTP relative to saline and assessed differences using an ANOVA model which specified current trial type (orienting versus inhibition), previous trial type (same versus switch), and number of preceding trials (1 versus 2) as fixed factors. We then examined performance separately for orientation and inhibition trials with separate ANOVA models specifying the previous trial type (same versus switch) and the number of preceding trials (1 versus 2) as fixed factors. Direct post hoc comparisons were made with two tailed independent *t*-tests and *P*-values were corrected for multiple comparisons with Tukey tests.

To examine if 5-HTP impacted how long monkeys took to initiate each trial, we quantified the time between the end of the inter-trial-interval and the next successfully initiated trial (i.e., acquiring the central fixation and maintaining the fixation for 150ms) during saline and 5-HTP sessions and compared them using a Wilcoxon rank sum test. To determine if the time taken to initiate trials was related to performance during each session, we correlated each session’s percent change in the time to initiate trials due to 5-HTP relative to saline and the percent change in performance due to 5-HTP relative to saline. For analyzing pupil size, we quantified the size of the pupil during the fixation period of initiated trials and compared pupil size between 5-HTP and saline sessions using a Wilcoxon rank sum test. We determined the relationship between pupil size and performance by correlating each session’s percent change in pupil size due to 5-HTP with the percent change in performance due to 5-HTP, both relative to saline. To examine the relationship between reaction time and performance, we first quantified reaction time as the time from cue/image onset to when animals began saccades to target images, using a 20deg/sec velocity criterion for detecting the saccade onset (i.e., reaction time). We assessed differences in reaction time during correct orientating trials using an ANOVA specifying animal identity (Monkey E, Monkey H, Monkey T), drug (saline versus 20mg/kg 5-HTP), image type (face versus scrambled), and image valence (threat versus appetitive) as fixed factors. We then correlated each session’s percent change in reaction time with the percent change in performance during orienting trials, both due to 5- HTP relative to saline. Direct post hoc comparisons were made with two tailed independent *t*-tests and *P*-values were corrected for multiple comparisons with Tukey tests. Correlations were reported by calculating a Pearson’s linear correlation coefficient.

## Results

Three rhesus macaques performed a task designed to test their ability to flexibly orient, or inhibit orienting, toward peripheral target images by using a colored instructional cue that appeared at the same time as the visual image (**Fig. 1A**). Peripheral target images were unfamiliar conspecific faces or luminance-matched scrambled faces as controls. We tested the causal role of serotonin in orienting and inhibiting gaze responses by increasing central concentration of serotonin with the direct precursor, 5-HTP. Our main behavioral measure of interest was the percent of trials correctly completed for both orienting and inhibition trials. We also investigated how the preceding behavioral context impacted orienting and inhibition performance. Finally, we examined the effect of 5-HTP on pupil size, how long monkeys took to initiate trials, and reaction time in order to relate these measures to changes in performance due to 5-HTP.

### 5-HTP increases central serotonin and constricts the pupil

We have previously demonstrated that 5-HTP increases concentrations of both 5-HTP and serotonin in cervical CSF (Weinberg-Wolf et al., 2018). In all three monkeys used in the study, CSF 5- HTP concentrations were higher after receiving 20mg/kg 5-HTP compared to saline (*P* < 0.01, *t*-test; **Fig. 1B**). Additionally, in all three monkeys, 5-HTP administration also increased central serotonin (*P* < 0.01; **Fig. 1B**). Importantly, central concentrations of 5-HTP and serotonin were strongly correlated with one another, indicating that increases in serotonin are proportional to increases in 5-HTP (*r* = 0.83, *P* = 0.02, Pearson’s correlation). Because cervical CSF taps are invasive procedures, we previously established a biomarker of 5-HTP’s impact on central serotonin; 5-HTP dose dependently decreases the size of the pupil (Weinberg-Wolf et al., 2018). We thus quantified the size of the pupil, following 5-HTP and saline administrations, during the 150ms fixation period where a luminance-controlled white fixation square appeared alone on the screen and when animal’s eyes were steady as they fixated. Monkeys exhibited a significantly more constricted pupil during 5-HTP sessions than saline sessions (*P* < 0.001, Wilcoxon sign rank; more later), and the magnitude of this constriction (−31.27 ± 1.18%, mean ± s.e.m.) was consistent with our previous work (Weinberg-Wolf et al., 2018). These cisternal CSF and pupil results support that 5-HTP administrations effectively cross the blood-brain barrier to causally influence central serotoninergic function and the parasympathetic system.

### Indifferent saccadic and fixation behaviors between 5-HTP and saline

To ensure that 5-HTP did not merely cause atypical gaze behaviors to influence performance, we compared saccade kinematics and gaze fixations during the task. The relationship between peak saccade velocity and amplitude did not differ between 5-HTP and saline sessions when monkeys oriented to faces (*P* = 0.94, permutation test) or to control images (*P* = 0.57) (**Fig. 1C**). We also compared gaze fixation patterns on target image locations during successfully completed orient trials, as well as gaze fixations on any parts of the monitor screen during correctly inhibited trials, between 5-HTP and saline sessions. Fixation patterns during orient trials were not different between 5-HTP and saline (the right and left visual image locations, both *P* > 0.06; Wilcoxon rank sum) (**Fig. 1D**). Moreover, monkeys largely remained fixated on the central cue during successful inhibit trials (despite the fact that the task did not require them to do so), and these fixation patterns again did not differ across 5-HTP and saline (*P* = 0.30) (**Fig. 1D**). Therefore, low-level gaze behaviors were not impacted by 5-HTP.

### 5-HTP impairs orienting and inhibition performance

Our primary question was whether increasing central concentrations of serotonin using 5-HTP would impact monkeys’ ability to orient or inhibit orienting toward faces. We first assessed performance by quantifying the percent of trials completed correctly over the course of a saline or 5-HTP session. We observed strong impairments in performance following 5-HTP (78.3 ± 1.7%, mean ± s.e.m.) compared to saline (85.9 ± 1.1%) (*F*(1, 336) = 75.34, *P* < 0.001, ANOVA; **Fig. 2A** and **2B**). We next queried how 5-HTP affected orienting and inhibition performance. Monkeys performed orienting trials at a near-ceiling performance after receiving saline (98.1 ± 0.2%), whereas they displayed orienting impairments after 5-HTP (90.8 ± 1.8%) (*P* < 0.001, Tukey test; **Fig. 2A** and **2B**). After normalizing performance to saline, we found that 5-HTP impaired orienting performance similarly on trials with control images (raw 5-HTP: 88.7 ± 2.8%, raw saline: 98.3 ± 0.3%, −5.3 ± 1.6% change from saline) and those with face stimuli (raw 5-HTP: 92.8 ± 2.2%, raw saline: 97.9 ± 0.4%, −9.6 ± 2.0% change from saline) (*F*(1,92) = 2.87, *P* = 0.09, ANOVA; **Fig. 2C**). We also asked if orienting performance differed according to the expressions conveyed by face images: threatening versus affiliative. However, monkeys oriented to threatening and affiliative faces at similar rates (*F*(1, 184) = 0.11, *P* = 0.74, ANOVA), and 5-HTP impaired performance during appetitive and threatening trials similarly (*F*(1, 92) = 0.04, *P* = 0.85). Although this negative finding may seem surprising at first, it was likely due to the fact that target images in this task disappeared as soon as monkeys’ gaze first entered the area, leaving no time for the monkeys to foveate on or scan them.

**Figure 2.**
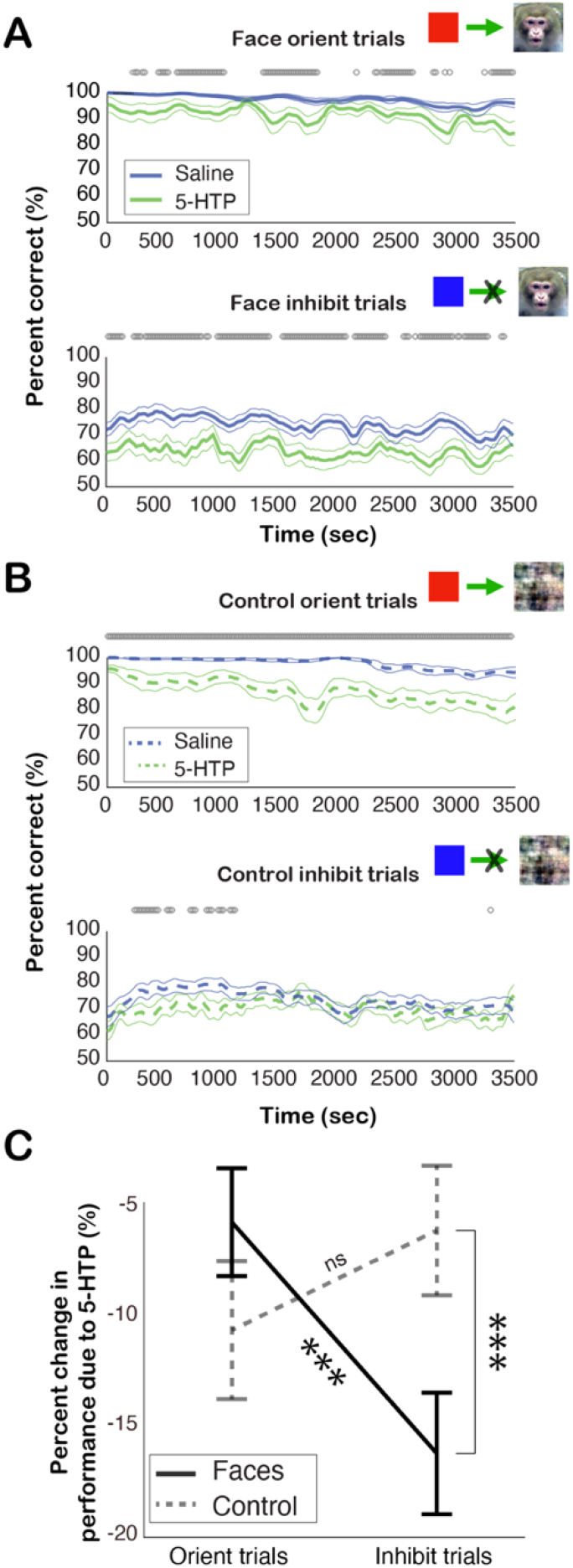
5-HTP disrupts orienting and inhibition performance. **A)** Average performance over the course of a session during orienting trials (top) and inhibition trials (bottom) for face images. **B)** Average performance over the course of a session during orienting trials (top) and inhibition trials (bottom) for scrambled control images. For the panels **A** and **B**, performance during saline sessions is shown in blue, whereas 5-HTP session is shown in green (mean ± s.e.m.). The grey circles above each time series represent significant differences between 5-HTP and saline at each time point (*P* < 0.05, Wilcoxon sign rank). **C)** The percent change in performance due to 5-HTP relative to saline (mean ± s.e.m.), illustrating that 5-HTP selectively impaired performance on inhibition trials with faces images (solid lines) but not those with control images (dashed line). ***, *P* < 0.001, ns, not significant, ANOVA.

We next investigated performance on inhibition trials. Monkeys were worse at performing inhibition trials (74.4 ± 1.5%, mean ± s.e.m.) compared to orienting trials (98.1 ± 0.2%) (*F*(1,184) = 24.99 *P* < 0.001, ANOVA) during saline sessions (**Fig. 2A** and **2B**), indicating that it was difficult for monkeys to inhibit looking at a stimulus in their periphery, and 5-HTP further decreased this performance (66.8 ± 1.5%, *P* < 0.001; Tukey test; **Fig. 2A** and **2B**). After normalizing performance to saline, we found that 5-HTP impaired inhibition performance more on trials with face images (raw 5- HTP: 66.8 ± 2.2%, mean ± s.e.m., raw saline: 74.7 ± 1.9%, −14.72 ± 1.9% change from saline) compared to those with control images (raw 5-HTP: 69.8 ± 1.9%, raw saline: 74.2 ± 2.3%, −5.36 ± 1.9% change from saline) (*F*(1,92) = 12.16, *P* < 0.0001, ANOVA; **Fig. 2C**). We again observed no differences in inhibition performance between trials with threatening compared to appetitive faces (*F*(1, 184) = 0.22, *P* = 0.64, ANOVA), and 5-HTP impaired performance during appetitive and threatening trials similarly (*F*(1, 92) = 0.44, *P* = 0.51). Overall, these measures suggest that increasing central serotonin availability with 5-HTP impairs orienting and inhibiting gaze responses, but that faces, perhaps due to their inherent saliency, caused an especial impairment in the ability to inhibit orienting gaze toward them (**Fig. 2C**).

### 5-HTP reduces flexibility in orienting and inhibiting gaze responses

Work across multiple species has investigated the role of serotonin in inhibiting responses and flexibly changing response (Roberts et al., 2020). In the current task, monkeys were trained to utilize a central cue to flexibly switch between two responses: orienting or inhibiting orienting. We tested flexibility in this context by considering the perseverating effect of preceding trials (Durston et al., 2002). To this end, we compared performance on trials where the previous trials (one or two) had been correct in a row as well as in which the response type (i.e., orient or inhibit) was either the same as, or a switch from, the current trial type in four distinct conditions (Materials and Methods).

After normalizing performance to saline, we found that 5-HTP disrupted performance more when the preceding trials were a switch from the current type (O→I, I→O, O→O→I, and I→I→O) compared to when they were the same (O→O, I→I, O→O→O, and I→I→I) (*F*(1,184) = 5.73, *P* = 0.02, ANOVA). Notably, we found that 5-HTP worsened performance on inhibition trials which were preceded by correct orientation trials (O→I and O→O→I versus I→I and I→I→I) (*F*(1,92) = 4.96, *P* = 0.03), but only when there had been two (O→O→I versus I→I→I, *P* = 0.03, Tukey test, **Fig. 3**) and not one (O→I versus I→I, *P* = 0.99) preceding trial. By contrast, 5-HTP did not alter orienting performance according to the previous type (I→O and O→O→I versus O→O and O→O→O) (*F*(1,92) = 1.11, *P* = 0.30) or previous trial sequence length (I→O and O→O versus O→O→I and O→O→O) (*F*(1,92) = 0.73, *P* = 0.39; **Fig. 3**).These findings suggest that 5-HTP increased the difficulty in inhibiting prepotent responses or, perhaps, that serotonin decreased the ability to flexibly switch between actions by increasing perseverance.

**Figure 3.**
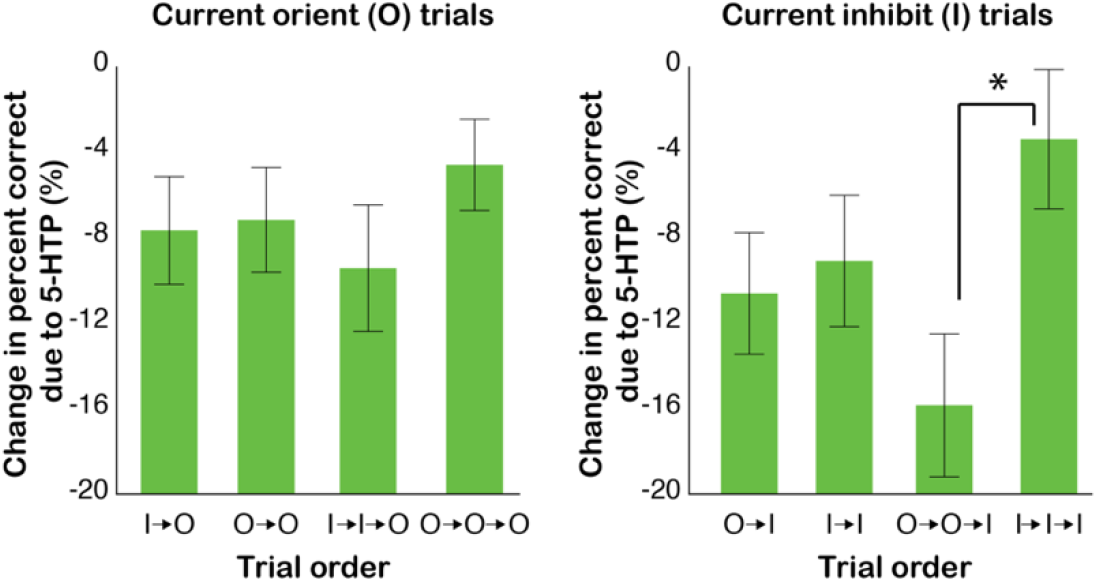
5-HTP reduces flexibility in orienting and inhibiting gaze responses in monkeys. 5-HTP impaired performance when the preceding trials were a swtich from the current trial type, but not when they were the same trial type (bars represent mean ± s.e.m.) *, *P* < 0.05, Tukey test. Trial order is shown as a sequence of response types (O, orientation or I, inhibition). For example, I→O indicates the current orientation trial followed a successfully completed inhibition trial in the previous trial. Similarly, O→O→I indicates the current inhibition trial followed a sequence of two successfully completed orientation trials.

### Decreases in task motivation and arousal by 5-HTP are related to performance impairments

Psychopathologies for which serotonergic interventions are relied upon, especially depression, are commonly associated with decreased motivation and anhedonia (Grahek et al., 2019). For this reason, we were curious if 5-HTP altered monkeys’ task engagement in the present task. To this end, we calculated the time it took for monkeys to successfully initiate a trial (Materials and Methods) based on the reasoning that under a more engaged state, monkeys ought to initiate trials more quickly compared to under a less engaged state. Monkeys took longer to initiate trials during 5-HTP sessions (5-HTP: 1800 ± 356ms, mean ± s.e.m., saline: 583 ± 50ms, *P* = 0.049, Wilcoxon rank sum; **Fig. 4A**, left). Crucially, after normalizing the number of initiated trials and performance of each session to saline, we found that the magnitude of 5-HTP’s effect on task engagement and performance were strongly, negatively, correlated with one another (*r* = −0.80, *P* < 0.001; **Fig. 4A**, right). Thus, 5-HTP concurrently increased the inter-trial initiation time and decreased performance on those trials.

**Figure 4.**
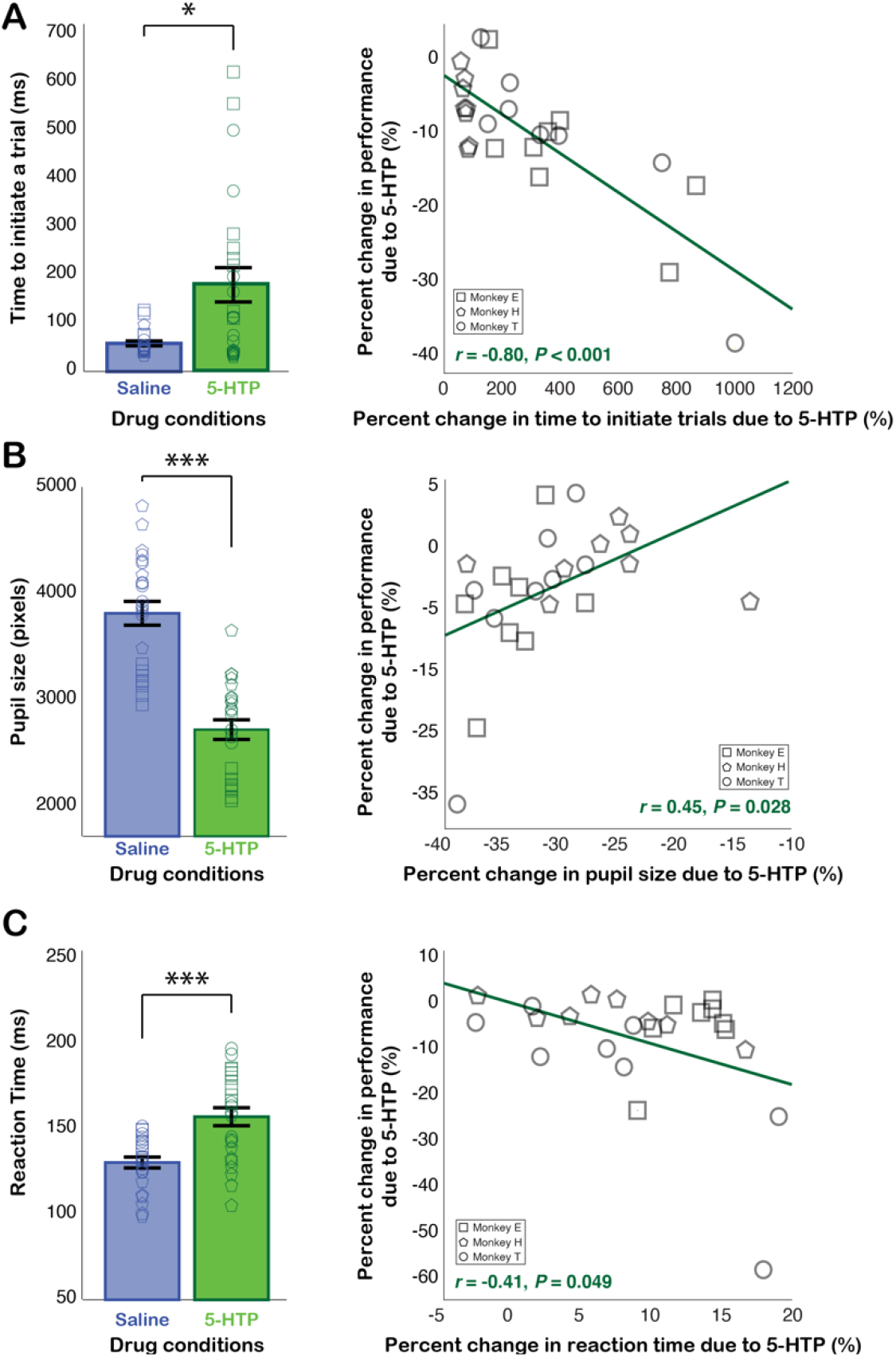
5-HTP increases inter-trial initiation time, constricts the pupil, and increases reaction time with concomitant changes in performance. **A)** (Left) 5-HTP increased the amount of time monkeys took before initiating a trial (mean ± s.e.m.; individual session data are overlaid with each monkey’s data using different shapes). (Right) The percent changes in the inter-trial initiation time were correlated with the percent changes in performance due to 5-HTP. **B)** (Left) 5-HTP constricted the pupil (same format as in the left panel of **A**). (Right) The percent changes in pupil constriction were correlated with the percent changes in performance due to 5-HTP. **C)** (Left) 5-HTP increased reaction time (same format as in the left panel of **A**). (Right) The percent changes in reaction time were correlated with the percent changes in orientation trial performance due to 5-HTP. Saline sessions are shown in blue, while 5-HTP sessions are shown in green. Each shape represents individual monkey’s data. *, *P* < 0.05, ***, *P* < 0.001, t-test. The green lines in the scatter plots in **A–C** illustrate linear regression fits.

Autonomic arousal, estimated by pupil size, is also related to motivation and engagement (Zekveld et al., 2014; Peysakhovich et al., 2015; van der Wel and van Steenbergen, 2018; Shechter and Share, 2020). Therefore, we were also interested in examining if potential changes to autonomic arousal, due to 5-HTP, influenced performance. To answer this question, we calculated the percent change in pupil size during the pre-trial fixation period (controlling for luminance and eye position) due to 5-HTP for each session and correlated it with the percent change in performance due to 5-HTP. We first quantified the size of the pupil following 5-HTP and saline administrations during the 150ms fixation period where a luminance-controlled white square appeared alone on the screen and when monkeys’ eyes were steady as they fixated. As mentioned earlier, 5-HTP constricted pupil compared to saline by nearly 31% (*P* < 0.0001, Wilcoxon sign rank; **Fig. 4B**, left). Notably, we observed a positive correlation between these two variables (*r* = 0.45, *P* = 0.028, Pearson’s correlation; **Fig. 4B**, right), indicating that the more 5-HTP constricted the pupil during a session, the more it also impaired performance. This finding further suggests that 5-HTP-induced impairments in task performance were linked to serotonin-mediated changes to autonomic arousal.

### 5-HTP concomitantly increases reaction time and impairs gaze orienting responses

Reaction time is commonly used to study attention (Prinzmetal et al., 2005) and motivation (Mir et al., 2011). We were curious as to whether increasing central concentrations of serotonin with 5-HTP would affect the reaction time of gaze orienting responses in the current task. Reaction time was longer during 5-HTP sessions (205 ± 3ms, mean ± s.e.m.) compared to saline sessions (188 ± 2ms) (*F*(1, 92) = 33.61, *P* < 0.001, ANOVA) (**Fig. 4C**, left). Reaction time did not, however, vary between trials with face and control images (*F*(1,92) = 0.21, *P* = 0.65) or threatening and appetitive faces (*F*(1,92) = 0.19, *P* = 0.67). We next queried the potential relationship between reaction time and performance during orientation trials by correlating the percent change in reaction time and the percent change in performance due to 5-HTP. Changes in reaction time were negatively correlated with changes in performance (*r* = −0.41, *P* = 0.049, Pearson’s correlation; **Fig. 4C**, right) such that the more 5-HTP increased reaction time in a given session, the more it also impaired orientation performance. This finding suggests that changes in reaction time may index a state change in attention, engagement, or motivational which is associated with 5-HTP’s impairments to orienting gaze responses.

## Discussion

Navigating one’s social environment requires individuals to flexibly orient to others, or inhibit orienting to them, depending on the context. While there are many benefits associated with collecting social information, there are also costs. Monitoring others not only causes animals to miss out on gaining food sources, but it also leaves them less time to monitor the environment for predators and environmental risks. There are also social risks associated with social gaze; direct eye contact can be considered a threat to rhesus macaques (Maestripieri, 1997), and macaques avoid the escalated aggression that can follow inappropriate social threats (Higley et al., 1996). Social gaze behaviors are therefore critical to competent social behavior, which is hypothesized to be associated with serotonergic function (reviewed in Weinberg-Wolf and Chang, 2019). In the current study, we tested how acute administrations of 5- HTP would alter macaques’ social orienting responses using gaze. 5-HTP selectively impaired social inhibition performance during trials with face images, whereas it impaired performance on all orienting trials similarly. While our scrambled images of faces were matched for low level characteristics like luminance, faces are still more salient and attentionally capturing (Shepherd et al., 2010). Therefore, faces might have captured attention more effectively and interacted with 5-HTP’s effects to disrupt inhibition performance more strongly.

Alternating between orienting and inhibiting responses requires flexibility. 5-HTP decreased performance on inhibition trials which were preceded by orienting trials, suggesting that 5-HTP induced monkeys to perseverate and become less flexible in switching between orienting and inhibiting gaze responses. Flexibility has traditionally been studied using reversal learning tasks and individual differences in flexibility have been related to serotonergic function (Barlow et al., 2015), serotonergic neurons (Matias et al., 2017), and differences in serotonergic related genetics (Izquierdo et al., 2007). In addition, casually depleting serotonin, particularly in the orbitofrontal cortex, impairs reversal learning (Clarke et al., 2004; Clarke et al., 2005; Walker et al., 2009) and increases stimulus stickiness (Rygula et al., 2015). Interestingly, a recent study has reported that oral 5-HTP impairs decision-making in an Iowa Gambling task and decreased the likelihood that subjects would switch choices between “decks” in the task (Gendle and Golding, 2010). While these effects do not seem to have been due to increased perseverance, they do indicate a reduction in flexible decision-making (Gendle and Golding, 2010). The serotonergic system is extremely complex with 14 serotonin receptor subtypes, each with differing functions (Roberts et al., 2020). For example, studies have shown that antagonizing the 5-HT2A receptor impairs performance in reversal learning tasks while antagonizing the 5-HT2C receptor enhances reversal performance (Boulougouris et al., 2008; Furr et al., 2012; Nilsson et al., 2012). Additionally, serotonergic drugs can also impact behavioral outcomes in a categorially different manner according to individual differences and differences in drug doses (reviewed in Weinberg-Wolf and Chang, 2019). Indeed it has been shown that while a small, acute, dose of the SSRI citalopram impairs reversal learning, a high, acute, dose had the opposite effect (Bari et al., 2010). Therefore, the impairment we observed in the current study, rather than improvement, in performance and flexibility is not wholly unexpected.

Disruptions to the serotonergic system have been casually linked to action impulsivity, especially through early responding in five-choice serial reaction time tasks and increases in firing rates of dorsal raphe nucleus neurons when rodents must wait for delayed rewards (Harrison et al., 1997b, a; Winstanley et al., 2005). Contrary to what one might predict from these results, increasing central serotonin with 5-HTP impaired inhibition performance in our task. It is worthwhile to note that our task was not designed to measure action impulsivity. While inhibition trials were more difficult for monkeys, the current task did not build a prepotent responses as observed in the tradition go/no-go or stop signal reaction time paradigms because orienting trials and inhibition trials occurred randomly, and at equal probability, and the target image appeared equally on the left and right side of the screen. Instead, our task required monkeys to use an instructional cue to select the appropriate gaze response.

The observed effects are likely mediated by serotonin’s effects on parasympathetic and autonomic arousal, as indexed by pupil size. Mental strain and focus increase the size of the pupil (Zekveld et al., 2014; Peysakhovich et al., 2015; van der Wel and van Steenbergen, 2018; Shechter and Share, 2020), while sleepiness and fatigue decreases pupil size (Hopstaken et al., 2015). Our lab has established that 5-HTP dose-dependently constricts the size of the pupil (Weinberg-Wolf et al., 2018), an effect we have replicated in the current study. This biomarker indicates that 5-HTP has a consistent effect on the parasympathetic nervous system and autonomic arousal. More specifically, we found that the change in pupil size was correlated with change in performance such that the more 5-HTP constricted the pupil during a session, the more it also impaired performance. In contrast to our results, SSRIs have been shown to dilate the pupil (McDougal and Gamlin, 2011), as does phasic activation of serotonergic neurons (Cazettes et al., 2020). Furthermore, serotonin syndrome, associated with pathologically high levels of serotonin, is also associated with pupil dilation (Alusik et al., 2014). However, early causal studies conducted directly with serotonin and 5-HTP reported causal pupil constriction (Reid and Rand, 1952; Page, 1954; Wada and McGeer, 1966; Rapport, 1997), suggesting potentially different downstream effects from elevating serotonin availability using 5-HTP versus through SSRIs. Our 5-HTP manipulations could be activating the parasympathetic nervous system, decreasing autonomic arousal, or a combination of the two. This effect on arousal may be critically underlying our observed changes in performance.

The observed effects of 5-HTP could also be due, at least in part, to changes to motivational state. Overall, the percent change in time monkeys took to initiate a new trial was negatively correlated with the percent change in performance such that the more 5-HTP increased the time to initiate a trial, the more it also impaired performance. This metric likely indexes motivational state and task engagement. Reaction time has also been used to estimate engagement (Mir et al., 2011). Here, the logic dictates that the more engaged animals are in a task, or the more motivated they are to view an image, the faster their reaction time would be. 5-HTP caused reaction times to slow with concomitant changes in performance, further supporting 5-HTP’s proposed role in affecting motivational state. It should be noted, however, that an increase in reaction time alone may not necessarily signify a decrease in engagement. For example, a previous study in humans have found that an acute dose of the SSRI citalopram increased participant’s response time and led to more careful moral decision-making (Crockett and Cools, 2015). Still, an earlier study investigated the effects of 30mg/kg 5-HTP on a bar pressing task in monkeys and found that 5-HTP transiently abolished bar pressing with concomitated pupil constriction, sleepiness, and decreased interest in favored foods (Wada and McGeer, 1966). We thus hypothesize that 5-HTP’s effects are primarily driven by a downregulation of arousal and motivational states, resulting in a constricted pupil, decreased number of trials initiated, slowing of reaction time, and impairment in performance.

Deficits in motivation are common in mood disorders (Grahek et al., 2019). While SSRIs are used to treat depression, they are also associated with indifference and decreased sexual motivation (Roberts et al., 2020). In addition, increased brain serotonin has been associated with decreased motivation to seek rewards (Roberts et al., 2020). However, SSRIs also sometimes increase motivation (Meyniel et al., 2016), although these effects could be due to inadvertent activation of the dopaminergic system (Subhan et al., 2000). Furthermore, amongst those at risk of depression, decreasing central serotonergic function with acute tryptophan depletion can decrease motivation (Roiser et al., 2006). The timeline of serotonergic interventions also seems to be critical. SSRIs often require chronic administration and can even cause a worsening of symptoms in patients at first (Duman et al., 2016; Roberts et al., 2020). In support of time-dependent effects of serotonin, optogenetically driving serotonergic cells was found to decrease acute, spontaneous, locomotion in mice, but repeated stimulation was found to increase spontaneous locomotion in the aggregate (Correia et al., 2017). Given the complex auto-receptors associated with the serotonergic system, acute and chronic manipulations may result in variable outcomes.

Overall, our findings provide causal evidence that acutely increasing central serotonin with the direct precursor 5-HTP impairs the ability to flexibly orient or inhibit orienting to faces, likely by affecting the motivational and arousal systems. Future work replicating these findings could further clarify the mechanism underlying these effects by directly testing 5-HTP’s effect on reversal learning, motivation, social attention, and engagement. In general, 5-HTP is highly understudied and the field would benefit from dissecting exactly how i.m. 5-HTP affects serotonergic receptor binding throughout the brain. Continued efforts toward better understanding the causal effects of 5-HTP on behaviors, circuits, and receptors would clarify the relationship between the serotonergic system and complex behaviors, and could therefore help improve matching treatment outcomes with patients.

## Acknowledgements

We thank Dr. Katalin Gothard for use of her image database. We also thank Dr. Amrita Nair for general assistance and Dr. George Anderson for quantifying the concentrations of serotonergic compounds in cerebrospinal fluid. The data used in the paper are available and downloadable through the GitHub page (https://github.com/changlabneuro/5HTP_orientinhibit.git).

